# Targeting of Protein Kinase CK2 Elicits Antiviral Activity on Bovine Coronavirus Infection

**DOI:** 10.1101/2021.06.08.447588

**Authors:** Ailyn C. Ramón, George V. Pérez, Evelin Caballero, Mauro Rosales, Daylén Aguilar, Dania Vázquez-Blomquist, Yassel Ramos, Arielis Rodríguez, Viviana Falcón, María P. Rodríguez, Yang Ke, Yasser Perera, Silvio E. Perea

**Affiliations:** Molecular Oncology Group, Department of Pharmaceuticals, Biomedical Research Division, Center for Genetic Engineering and Biotechnology (CIGB), Havana 10600, Cuba; Department of Animal and Human Biology, Faculty of Biology, University of Havana (UH), Havana 10400, Cuba; Pharmacogenomic Group, Department of System Biology, Biomedical Research Division, CIGB, Havana 10600, Cuba; Mass Spectrometry Laboratory, Proteomics Group, Department of Systems Biology, Biomedical Research Division, CIGB, Havana 10600, Cuba; Microscopy Laboratory, Department of System Biology, Biomedical Research Division, CIGB, Havana 10600, Cuba; Agricultural Research Department, Animal Biotechnology Division, CIGB, Havana 10600, Cuba; China-Cuba Biotechnology Joint Innovation Center (CCBJIC), Yongzhou Zhong Gu Biotechnology Co., Ltd, Lengshuitan District, Yongzhou City 425000, Hunan Province, China

**Keywords:** bovine coronavirus, protein kinase CK2, kinase inhibitor, CIGB-325

## Abstract

Coronaviruses constitute a global threat to human population since three highly pathogenic coronaviruses (SARS-CoV, MERS-CoV and SARS-CoV-2) have crossed species to cause severe human respiratory disease. Considering the worldwide emergency status due to the current COVID-19 pandemic, effective pan-coronavirus antiviral drugs are required to tackle the ongoing as well as future (re)emerging virus outbreaks. Protein kinase CK2 has been deemed a promising therapeutic target in COVID-19 supported by its *in vitro* pharmacologic inhibition and molecular studies on SARS-CoV-2 infected cells. CIGB-325 is a first-*in*-class synthetic peptide impairing the CK2-mediated signaling whose safety and clinical benefit have been evidenced in Covid-19 and cancer patients after intravenous administration. Here, we explored the putative antiviral effect of CIGB-325 over MDBK cells infected by bovine coronavirus (BCoV) Mebus. Importantly, CIGB-325 inhibited both the cytopathic effect and the number of plaques forming units with a half-inhibitory concentrations IC_50_ = 3.5 μM and 17.7 μM, respectively. Accordingly, viral protein accumulation at the cytoplasm was clearly reduced by treating BCoV-infected cells with CIGB-325 over time, as determined by immunocytochemistry. Of note, data from pull-down assay followed by western blot and/or mass spectrometry identification revealed physical interaction of CIGB-325 with nucleocapsid (N) protein and a *bona fide* cellular CK2 substrates. Functional enrichment and network analysis from the CIGB-325 interacting proteins indicated cytoskeleton reorganization and protein folding as the most represented biological processes disturbed by this anti-CK2 peptide. Altogether, our findings not only unveil the direct antiviral activity of CIGB-325 on coronavirus infection but also provide molecular clues underlying such effect. Also, our data reinforce the scientific rationality behind the pharmacologic inhibition of CK2 to treat coronavirus infections.

## 1. Introduction

Coronaviruses (CoVs) comprise a diverse group of enveloped positive-strand RNA viruses in the family Coronaviridae that infects a wide variety of hosts, including mammals and birds, and cause several respiratory and enteric diseases [1, 2]. These viruses comprise infectious bronchitis virus (IBV) in chickens, porcine epidemic diarrhea virus (PEDV), transmissible gastroenteritis virus (TGEV), swine acute diarrhea syndrome-CoV (SADS-CoV), bovine coronavirus (BCoV) as many others [3]. Four coronaviruses (HCoV-229E, HCoV-OC43, HCoV-HKU1, HCoV-NL63) are continuously circulating in the human population, causing common cold. However, three highly pathogenic coronaviruses (SARS-CoV, MERS-CoV and SARS-CoV-2) have crossed species to cause severe human respiratory disease [3]. In 2002, a human pathogenic CoV, severe acute respiratory syndrome (SARS-CoV) caused more than 8000 human infections in 2002–2003, with a case fatality of ~10% [4]. A decade later, a novel lethal zoonotic disease appeared in the Arabian Peninsula caused by the Middle East respiratory syndrome coronavirus (MERS-CoV) with a 36.1% mortality rate according to the World Health Organization (WHO) [5]. Finally, a novel coronavirus SARS-CoV-2 emerged from Wuhan, China in December 2019 and three months after became a pandemic disease with more than 150 million people infected and three million deaths across the world, therefore collapsing national health systems and strengthen the economic crisis worldwide (www.worldometers.info/coronavirus, accessed June 6, 2021).

In this context, vaccination represents a reasonable strategy to prevent or avoid coronavirus infection, however, vaccines fail, in part, due to virus resistance emergency [6]. The frequency and extension of the abovementioned epidemiological events highlight the urgent necessity of therapeutics solutions to treat viral infection. Taking into consideration the unprecedented outbreak of SARS-CoV-2 occurs in a scenery with no established active molecules against Betacoronavirus, the availability of effective pan-coronavirus antiviral drugs is required to tackle (re)emerging virus outbreaks. Considering the need of a prompt drug development, re-purposing of established drug with other therapeutics uses allow a fastest way to conduct the proof-of-concept in the clinic [7, 8].

Traditional antiviral strategies directed to viral targets often yield drug resistance; therefore, targeting relevant host factors involved in viral replication guarantee therapeutics with a wide-spectrum activity since families of viruses share common cellular signaling pathways and processes [9, 10]. Within host factors that viruses hijack during the infection process, kinases proteins play a fundamental role phosphorylating viral and host substrates [11]. Phosphorylation landscape during the SARS-CoV-2 life cycle revealed changes in the activity of 97 out of 518 human kinases [12]. Remarkably, protein kinase CK2 was found dramatically upregulated with a clear increase in phosphorylation of well-characterized substrates involved in the cytoskeleton organization mainly on the filopodia protrusions that promote virus egress and rapid cell-to-cell spread [12]. Consistently, an in vitro strong anti-SARS-CoV-2 activity was achieved with the CK2 specific inhibitor CX-4945 supporting the relevance of this kinase during the viral cycle [12].

CIGB-325 (formerly CIGB-300) is a synthetic peptide designed to target a subset of CK2 substrates by binding to the conserved phosphoacceptor sites and recently it has shown a direct impact over the CK2 enzymatic activity as classical inhibitors [13-15]. This biochemical feature supports the first-in-class attributes of CIGB-325 and entails important pharmacological differences compared with other available CK2 inhibitors. Clinical data have confirmed that CIGB-325 peptide is safe and well-tolerated when administered by intravenous infusion, whereas showed clinical efficacy in cancer patients [16, 17]. Of note, CIGB-325 has shown preliminary evidences of antiviral activity in different in vitro models. For instance, anti-HIV activity was reported as the peptide interferes with a putative NPM1-Rev interaction in cells and subsequently downregulates Rev-dependent gene expression [18]. Recently, a Phase I/II clinical trial with CIGB-325 showed benefits in Covid-19 patients with pneumonia with significant reduction of the pulmonary lesions and quick improvement of the clinical status at day 7 [19].

Some many aspects of disease caused by highly pathogenic human CoV can be recapitulated in animal CoV diseases. Therefore, animal coronavirus models are suitable and alternative systems for testing putative pan-coronavirus therapies [20]. Moreover, it appears as an advantage of such models the fact that does not require restrictive BSL-3 facilities. In this work we aimed to explore the antiviral activity of the anti-CK2 peptidic inhibitor CIGB-325 against coronavirus infection, using a bovine coronavirus (BCoV) model which belongs to the Betacoronavirus genus that includes murine hepatitis coronaviruses (MHV), porcine hemagglutinating encephalomyelitis virus (HEV), rat coronavirus (RtCoV), human respiratory coronavirus HCoV-OC43 and severe acute respiratory syndrome (SARS-CoV, SARS-CoV-2) [21]. BCoV causes enteric and respiratory disease in cattle, and is zoonotically transmissible among species, since BCoV-like viruses have been detected in wild ruminants and humans [22]. BCoV shares biological pathogenic and pneumoenteric properties with species of Coronavirus related to SARS-CoVs [23]. Thus, despite BCoV use a different cellular receptor to enter into the cells, replication, assembly and final egress of viral particles may share common viral and cellular factors [24, 25].

In this work, we tested for the first time the putative antiviral activity of CIGB-325 in Coronavirus infection in vitro using MDBK cells infected by BCoV. Besides, we unveil mechanistic clues supporting the antiviral activity. Our data also reinforce the rationale for pharmacologic inhibition of CK2 to treat diseases originated by these types of viruses.

## 2. Materials and Methods

### 2.1. Cell culture and BCoV Virus

Madin-Darby Bovine Kidney cell line (MDBK) (ATCC^®^ CCL-22™) was maintained in Dulbecco’s modified Eagle’s medium (DMEM) (Gibco, USA), supplemented with 10% inactivated fetal bovine serum (FBS) (Invitrogen, CA, USA) at 37 °C and 5% CO_2_. The BCoV (strain Mebus) was obtained from the Pasteur Institute in São Paulo, Brazil, and titrated in serial 1 log dilutions (from 1 log to 11 log) to obtain a 50% tissue culture infective dose (TCID50) on 96-well culture plates of MDBK cells. The plates were observed daily during 4 days for the presence of cytopathic effect (CPE) using an inverted optical microscope. The viral titer was calculated according to the Reed & Muench method based on eight replicates for titration.

### 2.2. Cell Cytotoxicity Assay and Drug Treatments

Cell cytotoxicity was determined using crystal violet staining. Briefly, 60 000 MDBK cells were seeded in flat-bottom 96-wells plates per well in DMEM medium with 10% fetal bovine serum (FBS), and a curve of serial dilutions (1:2) of CIGB-325 (3.12-200 μM) were added by triplicate. After 4 days, crystal violet stain was conducted as described in 2.4. The half-cytotoxic concentration CC_50_ was estimated from the fitted dose-response curves using the CalcuSyn software (Biosoft, Cambridge, United Kingdom).

### 2.3. Cytopathic Effect Assay

To evaluate the antiviral effect against BCoV, 260 000 MDBK cells were seeded in 24-well cell culture plates per well and incubated overnight at 37 °C in 5% CO_2_. After incubation, selected concentrations of CIGB-325, CX-4945, or F20-2 negative control peptide were added to cell monolayers for 1 h in serum-free DMEM. Interferon alpha-2b (IFN alpha-2b) at 500 IU/ mL was employed in this study as a positive control. Subsequently, 14 000 TCID50 of virus in 200 μL were added to each well of the plate. After 1h of incubation, the final volume was completed up to 1 ml and the appropriate drug concentration was maintained. Incubation was prolonged for 4 days at 37° C in 5% CO_2_ and the CPE was finally revealed by crystal violet staining. The antiviral activity rate was expressed as the drug concentration that protects 50% of CPE. The half-inhibitory concentration (IC_50_) values were estimated from the fitted dose-response curves using the CalcuSyn software (Biooft, Cambridge, United Kingdom).

### 2.4. Crystal Violet Assay

The crystal violet staining was used to evaluate cytotoxicity of CIGB-325 and verify viral cytopathic effect perceived by visual observation. After the incubation period, medium was removed from each well, crystal violet (1%) (Sigma, MO, USA) was added and incubated for 5 min at room temperature. After dye removal, wells were washed with water. Finally, absorbance at 578 nm was read using a CLARIOstar^®^ high-performance monochromator multimode microplate reader (BMG LABTECH, Ortenberg, Germany).

### 2.5. Plaque Reduction Assay

Viral titers in supernatants of drug-untreated and treated cells were evaluated using a plaque reduction assay. Briefly, MDBK cells were seeded at 260 000 cells per well in 24-well cell culture plates and incubated overnight at 37 °C in 5% CO_2_. Subsequently, 200 μL of supernatants in 10-fold serial dilution was added to the MDBK monolayers. After 1 h of incubation at 37°C and 5% CO_2_, the viral inoculum was aspirated, and 0.5 mL of carboxymethylcellulose (Sigma, MO, USA) overlay with DMEM and supplemented with 2% FBS, was added to each well. After 4 days of incubation, the cells were fixed and stained with Naphtol Blue Black (0.1%) (Sigma, MO, USA MO, USA). Finally, plaques were counted visually and the virus titer as plaque-forming units (PFU) per mL was calculated.

### 2.6. Quantitative PCR Assay

MDBK cells were plated on 24-well cell culture plates at 260 000 cells per well and incubated overnight at 37 °C and 5% CO_2_. Once cell monolayers were established, CIGB-325 (15 μM, 30 μM), F20.2 (30 μM), CX-4945 (1.25 μM) or vehicle (PBS) were added for 1 h in serum-free DMEM. Subsequently, 14 000 TCID50 of virus in 200 μL were added to each well. After 1h of incubation, final volume was completed up to 1 ml and the appropriate drug’s concentration was maintained for 24 h. Three replicates per condition were used. After incubation time, the culture medium was withdrawn and the cells were washed with PBS and suspended in 350 μL of Lysis Buffer (with 1% of β-mercaptoethanol, Sigma, MO, USA) for RNA isolation (AllPrep DNA/RNA/miRNA Universal Kit, Qiagen, USA) according to the manufacturer protocol. All RNA samples were checked by Nanodrop spectrophotometer to measure concentration (ng/μL) and OD relation (260/280 nm). Quality control parameters were fulfilled by all the samples (100% OD 260/280 nm between 1.7 and 2.2). Complementary (c)DNAs were obtained from 500 ng of total RNAs, using the Transcriptor First Strand cDNA Synthesis Kit package (Roche, Germany) following manufacturer instructions. The qPCR reactions were set up in 20 μL using LightCycler^®^ 480 SYBR Green I Master 2x (Roche, Germany), 300 nM of oligonucleotides and 1:10 dilutions of each cDNA with three replicates per sample. We amplified two genes for normalization *gapdh* (Glyceraldehyde-3-phosphate dehydrogenase) and *hmbs* (Hydroxymethyl-bilane synthase) and the transcript encoding for N protein from BCoV. In parallel, we used a plasmid with the BCoV N protein transcript cloned that it was amplified from serial dilutions as a standard curve. This standard curve was used for N copy number calculations. All the oligonucleotides were synthetized in the Synthesis Department at CIGB (Table S1). Runs were carried out in the LightCycler^®^480II equipment (Roche, Germany) in a 96-well format and SYBR Green Probe II mode with a standard program with 45 cycles. Fold changes of N transcript expression with each treatment were calculated corcenning to the Virus control using REST 2009 program (v2.0.13, Qiagen, GbmH) [26], after normalization with *gapdh* and *hmbs* genes. using Ct values and reaction efficiencies per amplicon calculated in LinReg 2009 (v 11.3) [27]. Statistical differences were reported in this program with a p value associated significant for p < 0.05 [28]. Additionally, we also report a copy number of N transcript by extrapolation of Ct values into the regression formula from the standard curve (N-BCoV encoding plasmid copy number *vs* Ct).

### 2.7. BCoV Viral Proteins Detection by Western Blot

MDBK cells were plated on 24-well cell culture plates at 260 000 cells per well and incubated overnight at 37 °C in 5 % CO_2_. Once cell monolayers were established, CIGB-325 (30 μM) or vehicle (PBS) were added for 1h in serum-free DMEM. Subsequently, 14000 TCID50 of virus in 200 μL were added to each well of the plate. After 1h of incubation, final volume was completed up to 1 ml and the appropriate drug’s concentration was maintained for 24 h. After incubation, the culture medium was withdrawn and the cells were washed with PBS and lysed in RIPA buffer containing protease/phosphatase inhibitor (Thermo Fisher Scientific, MA, USA). Equal amounts of protein (30 μg/sample) were resolved in 12% SDS-PAGE. Proteins were then transferred to nitrocellulose membrane and immunoblotted with human polyclonal IgG anti-SARS-CoV-2 generated by CGIB and validated by ELISA and western blot. Detection was performed with peroxidase-conjugated anti-human IgG 1:100 (Jackson ImmunoResearch, USA) and signal was developed using SuperSignal West Pico Chemiluminescent Substrate (Thermo Fisher Scientific, MA, USA).

### 2.8. BCoV Viral Proteins Detection by Immunocytochemistry

MDBK cells were plated on eight well glass slide and incubated overnight at 37 °C and 5% CO_2_. After incubation, cells were pre-treated for 1 h with CIGB-325 (30 μM) or vehicle (PBS) and infected with 100 μL of virus at a concentration of 70 000 TCID50/well. After 1h of incubation, final volume was completed up to 500 μL and the appropriate drug’s concentration was maintained for 16 h and 24 h. Subsequently, the cells were washed with cold PBS three times and fixed in 4% formaldehyde for 10 min at 4 °C. After permeabilization with 0.5% Triton X-100 for 10 min, cells were blocked by incubation with 4% bovine serum albumin (Sigma, MO,USA) in PBS for 30 min at room temperature, washed again, and incubated with 30 μg/mL human polyclonal anti-SARS-CoV-2 (CIGB, Cuba) for 2 h at room temperature. Then, peroxidase-conjugated anti-rabbit secondary antibody 1:100 (Sigma, MO, USA) was incubated for 1 h at room temperature and washed 3 times with PBS. Finally, AEC substrate (Abcam, Cambridge, United Kingdom) was added and coverglass was mounted using 40% glycerol mounting medium and analyzed using a BX43 upright microscope (Olympus, USA).

### 2.9. Pull-down Assay

CIGB-325 interacting proteins were identified by *in vitro* pull-down followed by tandem mass spectrometric analysis (LC-MS/MS). MDBK cells were seeded at 3600000 cells in 60 mm dishes and infected with 500 μL of virus at 70,000 TCID50/dish. After 48 h post-infection, cells were collected, washed and lysed in hypotonic PBS solution (1×) (Sigma, MO, USA) containing 1 mM DTT (Sigma, MO, USA), TritonX-100 (1%) and complete protease inhibitor (Roche, Basel, Switzerland). Cellular lysates were cleared by centrifugation and 300 μg of total protein were incubated with biotin-tagged CIGB-325 (100 μM) or medium alone for 30 min at room temperature and added to 30 μL of pre-equilibrated streptavidin-sepharose matrix (GE Healthcare, IL, USA). Following 1 h at 4 °C, the matrix was collected by centrifugation and extensively washed with cold PBS 1 mM with DTT. CIGB-325 interacting proteins were eluted, resolved in an SDS-PAGE gel and processed as described later. For *in vivo* pull-down assays, BCoV-Mebus infected and uninfected MDBK cells were treated with biotin-tagged CIGB-325 (100 μM) or medium alone for 30 min at 37° C in 5% CO_2_. Subsequently, cells were collected and pull-down assay was conducted as above mentioned. Proteins bound to streptavidin-sepharose were eluted, resolved in a 12 % SDS-PAGE gel, and transferred onto nitrocellulose membranes. For western blot analysis, a human polyclonal IgG anti-SARS-CoV-2 (CIGB, Cuba) and a mouse monoclonal antibody against CK2α (Abcam, Cambridge, United Kingdom) were used as primary antibodies. Detection was performed with peroxidase-conjugated anti-mouse IgG 1:5000 (Sigma, MO, USA) and anti-human IgG 1:100 (Jackson ImmunoResearch, USA). In parallel, untreated cells were subjected to the same experimental procedure to identify those proteins non-specifically bound to streptavidin-sepharose matrix.

### 2.10. LC-MS/MS Analysis and Protein Identification

Proteins bound to streptavidin-sepharose matrix of both, CIGB325 treated sample and background control were processed by the FASP method using Microcon 30k centrifugal ultrafiltration units (Merck, Darmstadt) [29]. A volume corresponding to 4 μg of the peptide mixture was separated by a Proxeon Easy-nLC System (Thermo Fisher Scientific) using an RP-C18-A2 column (3 μm), 75 μm i.d. × 10 mm (Thermo Fisher Scientific) connected online to a hybrid quadrupole orthogonal acceleration tandem mass spectrometer QTof-2 (Water, MA, USA). The Masslynx system (version 3.5) from Water (MA, USA) was used for data acquisition and processing. Protein identification based on MS/MS spectra was made against the Swiss-Prot database by using Mascot database search engine (version 2.5). False discovery rate (FDR) was set to 1% for peptide and protein identification. Proteins identified uniquely in the CIGB-325 treated sample were considered as CIGB325 interacting proteins. Protein-protein interaction (PPI) networks were generated using information retrieved form STRING database [30], and protein kinase CK2 substrates among CIGB-325 interacting proteins were identified based on Meggio and Pinna dataset [31], the list of high confidence CK2 substrates reported by Bian et al. [32] and the PhosphoSitePlus database (www.phosphosite.org, accessed June 6, 2021).

### 2.11. Confocal Microscopy

MDBK cells were plated on 8-well glass slide and incubated overnight at 37° C and 5% CO_2_. After incubation, cells were infected with 100 μL of virus at a concentration of 70 000 TCID50/well. Uninfected cells were established as a negative control of the experiment. After 48 h post-infection, cells were treated with fluorescein-tagged CIGB-325 (30 μM) or vehicle (PBS) for 30 min at 37° C and 5% CO_2_. Subsequently, cells were washed with cold PBS three times and fixed in 10% formalin for 10 min at 4 °C. After permeabilization with 0.2% Triton X-100 for 10 min, cells were blocked by incubation with 4% bovine serum albumin (Sigma, MO, USA) in PBS for 30 min at room temperature, washed again, and further incubated with 30 μg/mL rabbit polyclonal anti-N protein for 2 h at room temperature. Finally, anti-rabbit IgG Alexa Fluor 594 conjugate at 1:250 dilution (Cell Signaling Technology, MA, USA) was incubated for 1 h at room temperature and washed 3 times with PBS. Coverglasses were mounted using Vectashield mounting medium with DAPI (Vector Laboratories, CA, USA) and analyzed using an Olympus FV1000 confocal laser scanning microscopeIX81 laser scanning fluorescence microscope (Olympus, Tokyo, Japan). Images were acquired with UPLSAPO 40× immersion objective and processed using Olympus FluoView software (v4.0) (Olympus, Tokyo, Japan). Five optical fields or Z-stacks were examined for each experimental condition.

### 2.12. CK2 Signaling Experiments

MDBK cells were plated on 24-well cell culture plates at 260 000 cells per well and incubated overnight at 37° C and 5% CO2. Once cell monolayers were established, 14 000 TCID50 of virus in 200 μL were added to each well of the plate. After 1 h of incubation, final volume was completed up to 1 mL of serum-free DMEM and incubated. After 24 h post-infection, cells were treated with CIGB-325 (30 μM) or vehicle (PBS) in serum-free DMEM for 45 min at 37 °C in 5% CO_2_. Subsequently, the culture medium was withdrawn and the cells were washed with PBS and western blot was carried out as described (2.7). Primary antibodies against phospho-RPS6 (S235/236) and total RPS6 (Cell Signaling Technology, MA, USA) were used, and detection was performed with peroxidase-conjugated anti-rabbit IgG 1:5000 (Sigma, MO, USA).

### 2.13. Statistical Analysis

Differences between groups were determined using one-way ANOVA followed by Dunnett’s multiple comparisons test. Analyses were performed in GraphPad Prism (v6.01) software for Windows (GraphPad Software, Inc, San Diego, CA, USA). All experiments were conducted at least in triplicates and differences were considered significant for a *p*-value < 0.05.

## 3. Results

### 3.1 CIGB-325 Exhibits Antiviral Effect on Bovine Coronavirus Infected Cells

To test the *in vitro* CIGB-325 effect, we analyzed the CPE of BCoV-Mebus using a cell-based assay. MDBK cells appeared as rounded and multinucleated cells within 48 h; beyond 72 h of incubation a strong cell monolayer damage was observed. CIGB-325 elicited a very potent inhibition of the virus CPE with an IC_50_=3.5 μM just some significant cytotoxicity emerges at higher concentrations (CC_50_=150μM) (**Figure 1A**), hence indicating a selectivity index (SI=42). Specificity of the CIGB-325 antiviral effect was corroborated using the F20-2 peptide which contains the same cell-penetrating peptide as CIGB-325 but lacks the CK2 inhibitory domain [13]. Interestingly, testing of another small molecule inhibiting CK2 activity (CX-4945), also reduced significantly the BCoV-Mebus CPE as well as IFN alpha-2b, which was used as an antiviral drug reference (**Figure 1B**).

**Figure 1.**
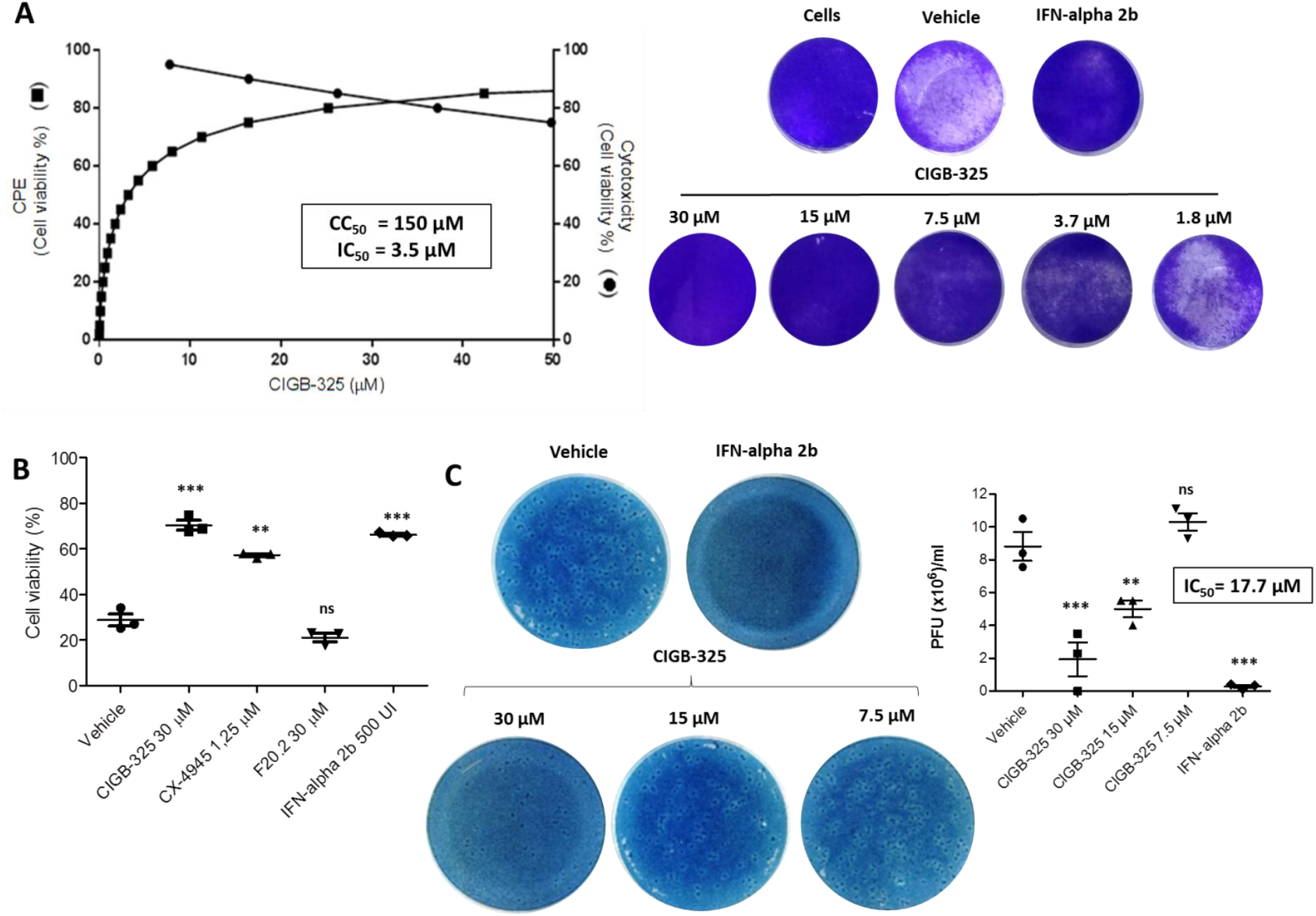
*In vitro* antiviral activity of CIGB-325 against BCoV-Mebus. MDBK cells were pre-treated with CIGB-325 at the indicated doses for 1 h, and virus (14 000 TCID50/well) was then added to allow attachment for 1 h. Afterward, the cells were incubated in the presence of the indicated drug concentration for 4 days. **A.** IC_50_ (left axis) and CC_50_ (right axis) were estimated from the fitted dose-response curves based on treatment with five CIGB-325 concentrations determined by cell viability assay (left section). Crystal violet stain wells for each experimental condition in the CPE inhibition assay (right section). **B.** Antiviral effect of positive (IFN-alpha2b, CX-4945) and negative controls (F20.2) were determined by cell viability assay using crystal violet stain. Cells were infected as previously described and incubated with CIGB-325 (30 μM), F20.2 (30 μM), CX-4945 (1.25 μM) and IFN-alpha2b (500 IU). Uninfected cells with the vehicle were set to 100%. **C.** Progeny titer of the cell cultures supernatants collected from CPE inhibition experiments were assessed using plaque assay. Data from **B.** and **C.** are shown as mean ± SD, n = 3. Statistically significant differences between vehicle and drug treatment are represented as \*\**p* < 0.01 and *** *p* < 0.001 determined using one-way ANOVA followed Dunnett post-test.

To corroborate the CIGB-325 inhibitory effects over BCoV-Mebus infection, we additionally measured the viral load on cell supernatants collected from CPE experiments by performing a plaque assay. CIGB-325 displayed dose-dependent inhibition of viral titers CIGB-325 according to the number of BCoV-Mebus PFU (**Figure 1C**). Of note, the CIGB-325 antiviral effect was achieved even 16 and 26 h post-viral challenge indicating a possible impact on the late stages of the BCoV-Mebus life cycle (**Figure S1**).

### 3.2 CIGB-325 Impacts the BCoV-Mebus Viral Machinery in MDBK Cells

To assess the putative effect of CIGB-325 over viral mRNA expression we conducted quantitative real-time PCR on BCoV-Mebus infected cells. Data from **Figure 2A** indicates that CIGB-325 significantly downregulated the viral nucleocapsid (N) protein copy number after 24 h of incubation in a dose-dependent manner while F20-2 slightly. Likewise, CX-4945 treatment affected the nucleocapsid gene expression, thus confirming a direct involvement of the CK2-mediated phosphorylation into coronavirus gene expression.

**Figure 2.**
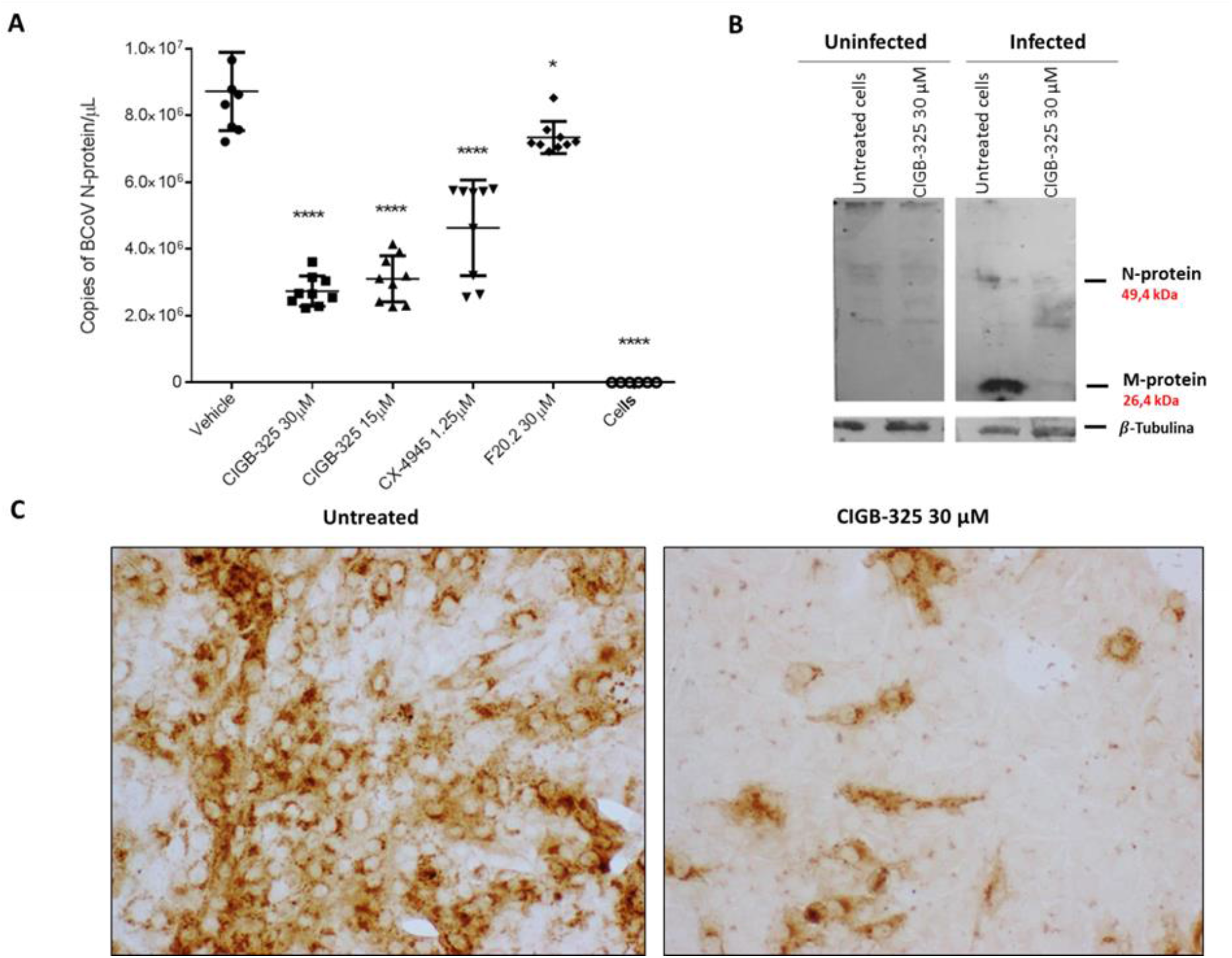
Impact of CIGB-325 over BCoV-Mebus viral machinery. **A.** MDBK cells were treated with the drugs 1 h before the viral challenge. Afterward, cells were infected with BCoV-Mebus (70 000 TCID50/well) and the appropriate drug’s concentration was maintained. Effect of CIGB-325 over the viral protein expression at 24 h post-infection was investigated using qRT-PCR (**A**), Western blot (**B**) and Immunocytochemistry (**C**). qRT-PCR reaction was performed with specific oligonucleotides corresponding to N protein. N expression was normalized to *gapdh* and *hmbs* genes, and expressed as copy number obtained after extrapolation in a calibration curve. Values are presented as mean ± SD (n = 9). Statistically significant differences between vehicle and drug treatment are represented as \*\**p* < 0.01 and \*\*\**p* < 0.001 determined using one-way ANOVA followed Dunnett post-test. Cell preparations for western blot and Immunocytochemistry were carried out as described in Materials and Methods. A human anti-SARS-CoV-2 polyclonal antibody from one COVID-19 convalescent patient was used for identification of BCoV-Mebus M and N protein. Anti-β-tubulin blot was used as loading control. Viral proteins and β-tubulin protein were blotted in the same gel.

Aside from viral mRNA levels, we also investigated the effect of CIGB-325 over viral protein synthesis. Western blot experiments were conducted using a human anti-SARS-CoV-2 polyclonal antibody from one COVID-19 convalescent patient. Of note, BCoV-Mebus M and N proteins were the most efficiently recognized by the human polyclonal antibody, exhibiting molecular weights of 49 and 26 kDa, respectively.

CIGB-325 treatment dramatically downregulated the viral M-protein expression at 24 h post-BCoV-Mebus infection, although the N protein levels were also affected to a lesser extent (**Figure 2B**). Immunocytochemistry data revealed that CIGB-325 treatment notably cleared the intracellular accumulation of both viral proteins at 24 h post-infection (**Figure 2C**). Likewise, as early as 16 h, CIGB-325 treatment also downregulated the intracellular viral proteins (**Figure S2**).

Considering that CIGB-325 exerts anti-CK2 activity by direct binding to CK2 conserved phosphoacceptor domain on substrates [13], we searched for such aminoacidic motifs on viral proteins to anticipate putative physical interactions. As we confirmed that BCoV-Mebus viral N protein displays such a sequence [33], we conducted *in vitro* pull-down experiments using biotinylated CIGB-325 to capture interacting proteins from infected-cell lysates with further binding to a streptavidin-sepharose matrix. Interestingly, western blot data from pull-down fractions using a human anti-SARS-CoV-2 polyclonal antibody showed that pull-down fractions but not the input, displayed a unique band corresponding to BCoV-Mebus N protein (49 kDa) (**Figure 3A**). Furthermore, *in vivo* pull-down assays were performed on BCoV-Mebus infected MDBK cells to confirm the interaction of CIGB-325 with viral N protein on a relevant cellular context. Of note, a clear band corresponding to the N protein (49 kDa) was again observed after western blot analysis (**Figure 3A**). To verify the *in situ* physical proximity between CIGB-325 and N protein at the subcellular compartments, we next employed confocal microscopy on infected MDBK cells treated with CIGB-325-F. After 30 min of incubation, a quite clear orange co-localization pattern of CIGB-325 with the viral N protein was observed mainly throughout the cytoplasm, with a slighter extent within the nucleus of the cell (**Figure 3B**).

**Figure 3.**
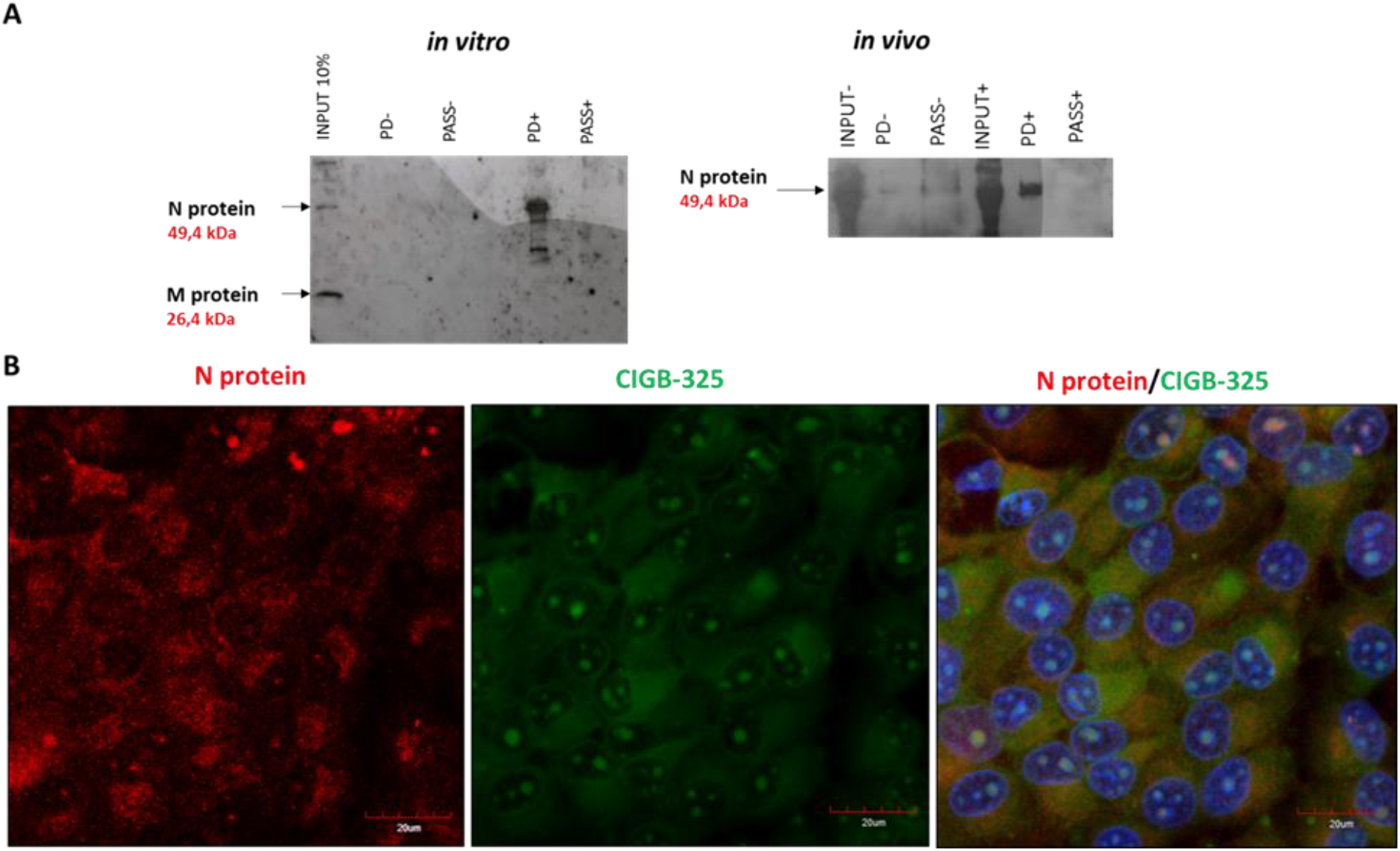
Interaction of CIGB-325 with BCoV-Mebus N protein. **A.** Western blot analysis of *in vitro* and *in vivo* pull-down fractions using CIGB-325 conjugated to biotin as bait to capture interacting proteins. *In vitro* pull-down was performed with cellular lysates from infected MDBK cells incubated 30 min with biotin-tagged CIGB-325 (100 μM). Subsequently, 30 μL of streptavidin-sepharose were added to each reaction and the CIGB-325 interacting proteins were eluted, resolved in 12%-SDS-PAGE and subjected to western blot to identify the viral N protein. For *in vivo* pull-down, MDBK cells were treated 30 min with biotin-tagged CIGB-325 (100 μM), subsequently lysed and processed as indicated above. PD: pull-down fractions; PASS: flow-through fractions; NC: negative control (cellular lysate from MDBK cells incubated with the vehicle). **B.** Representative images obtained by confocal microscopy showing the co-localization of CIGB-325-F with BCoV-Mebus N protein after 30 min of incubation on infected MDBK cells. Red fluorescence: N protein-derived signal; Green: CIGB-325-derived signal; Blue: nuclear DAPI; Orange: merge of Green/Red channels representing co-localization signal.

Previous findings have indicated that SARS-CoV-2 N protein directly interacts with protein kinase CK2 [34]. To corroborate such interaction in our *in vitro* BCoV-Mebus infection model, we conducted immunoprecipitation assays from infected MDBK cell lysates using a commercial anti-CK2 antibody and subsequent western blot with a rabbit polyclonal antibody against the viral N protein. Similarly, data revealed that CK2 also associates physically to the BCoV-Mebus N protein in this cellular context (**Figure S3**).

### 3.3 CIGB-325 Interactomic Landscape on BCoV-Mebus Infected MDBK Cells

To identify the full array of viral and host proteins which interact with CIGB-325 in BCoV-Mebus infected MDBK cells, we conducted *in vitro* pull-down experiments coupled to LC-MS/MS analysis. Lysates from MDKB treated or not with biotinylated CIGB-325 were separated by SDS-PAGE. After tryptic digestion, the resulting peptides were isotopically labeled, pooled together and analyzed by LC-MS/MS. Using this experimental approach, we identified 50 host proteins and 2 viral proteins that interact with CIGB-325 in infected MDBK cells. Viral proteins corresponded to N protein and Non-structural protein 2a (NSp2a). Members of the family of proteins including actin, tubulin, myosin, heat shock protein 70 (HSP70), heat shock protein 90 (HSP90), tyrosine 3-monooxygenase/tryptophan 5-monooxygenase activation protein as well as others showed in **Table S2** were identified as host CIGB-325 interacting proteins. In agreement with CIGB-325 interactome from tumor cells [14, 35], nucleolar protein B23/NPM1 was also identified in our study.

To envisage the putative biological processes that might be perturbed by CK2 targeting, enrichment analysis of the CIGB-325 interacting host proteins was performed. Of note, protein folding and the response to unfolded protein, cytoskeleton organization and cell cycle were significantly represented above all biological processes in the interactomic profile (**Figure S4**). Furthermore, we searched for *bona fide* CK2 substrates among the CIGB-325 interacting proteins according to Meggio and Pinna dataset [31], the list of high confidence CK2 substrates reported by Bian et al. [32] and the PhosphoSitePlus database (www.phosphosite.org, accesed June 6, 2021). Remarkably, CIGB-325 interacted with 15 proteins previously described as CK2 substrates in MDBK cells (**Table S2**). To better understand the global putative connection among the CIGB-325 interactome on infected MDBK cells, we constructed a protein-protein interaction (PPI) network with the identified proteins using information annotated in STRING database (**Figure 4A**). In PPI network, we detected two functional physiological complexes corresponding to proteins involved in cytoskeleton reorganization and protein folding, consistently with the most represented biological processes.

**Figure 4.**
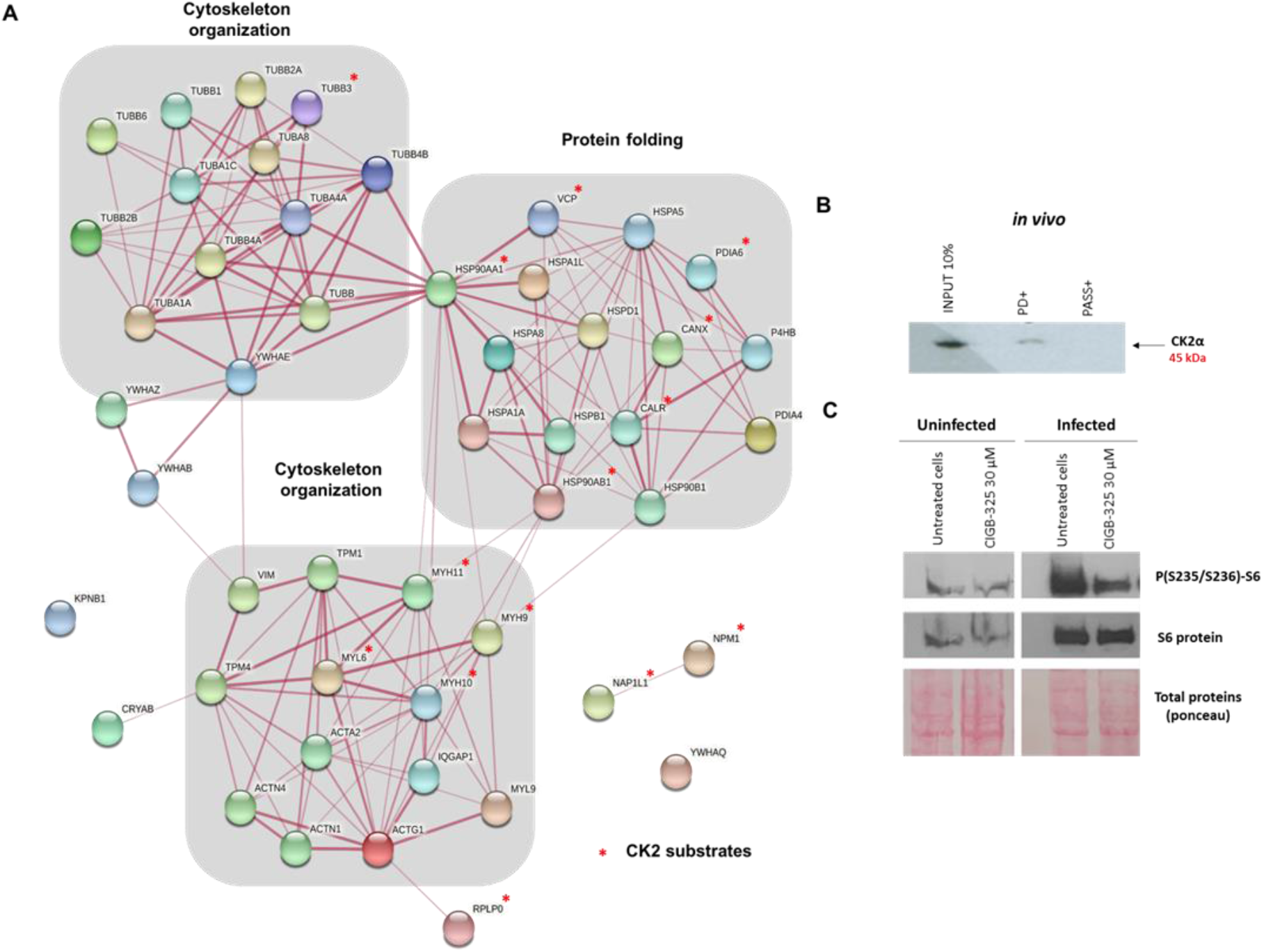
Protein-protein interaction network associated with CIGB-325 host interacting proteins on infected MDBK cells and subsequent effect over CK2. **A** Network was generated using information gathered from STRING database. Biological processes retrieved from Gene Ontology database are indicated and CK2 *bona fide* substrates are indicated with an asterisk in red. **B.** *In vivo* pull-down was performed with biotinylated CIGB-325 as bait to capture CK2 in MDBK infected cells. Interacting proteins were then resolved by 12% SDS-PAGE, transferred, and each fraction was inspected by western blot with the antibody against CK2α. **C.** Effect of CIGB-325 over phosphorylated and total protein levels of RPS6. MDBK cells were infected with BCoV-Mebus (70 000 TCID50/well). After 24 h of incubation, CIGB-325 was incubated for 45 min and preparation of cell extracts for western blot was carried out as referred to in Material and Methods.

Previous reports from our group have indicated that CIGB-325 also impairs CK2 signaling by direct binding to CK2α catalytic subunit in tumor cells [14, 15]. Therefore, we finally move forward to determine if this direct peptide-enzyme interaction could also take place within the context of a viral infection induced by BCoV-Mebus in MDBK cells. Using *in vivo* pull-down with biotinylated-CIGB-325 followed by western blot with a commercial anti-CK2α monoclonal antibody, it was possible to demonstrate the presence of CK2α in the pull-down fraction (**Figure 4B**). To corroborate whether this interaction could impact any of the CK2-mediated signaling pathways, we also evaluated the effect of CIGB-325 over the PI3K/Akt pathway by measuring the downstream phosphorylation of the RPS6 protein, as biomarker of the PI3K/Akt pathway activation which in turns, represents one of the CK2 mediated signaling pathways. Importantly, CIGB-325 treatment during 45 min clearly impaired the RPS6 phosphorylation (i.e.; at residues S235/236) at 24 h post-BCoV infection on MDBK cells (**Figure 4C**). However, a significant increase in RPS6 phosphorylation was observed upon BCoV-Mebus infection in these cells which reinforces the relevance of this pathway for the viral replication processes in the coronavirus biology [36, 37].

## 4. Discussion

During the last 20 years, three zoonotic CoVs, SARS-CoV, MERS-CoV and SARS-CoV-2 have evidenced that novel highly pathogenic coronavirus could spread into human populations [38]. Therefore, the development of an effective antiviral drug against already-known and new coronaviruses that may emerge from animal reservoir hosts constitutes an urgent necessity. Host-directed drugs represent a therapeutic strategy with broad-spectrum activity since viruses commonly hijack cellular factors and signaling pathways for their own replication [9, 39]. Among the host proteins relevant on some steps of the viral life cycle, protein kinase CK2 has been recently suggested as a fundamental factor in the SARS-CoV-2 infection [12]. Several evidences have shown an up-regulation of this protein during the viral replication along with a relevant molecular interaction with SARS-CoV-2 N protein at the filopodia protrusions. In fact, the specific CK2 inhibitor CX-4945 has displayed *in vitro* antiviral activity against SARS-CoV-2 infection [12]. Of note, the CK2-N protein interaction is conserved among the three coronaviruses (SARS-CoV, MERS-CoV and SARS-CoV-2) [10], therefore the scientific rationality of using CK2 inhibitor on pan-coronavirus infection could provide clinical benefit for future outbreaks.

CIGB-325 is a peptide-based drug capable to inhibit the CK2-mediated phosphorylation on the substrates, from clinic evidences previously assessed in cancer patients, this anti-CK2 peptide is safe and well-tolerated [16, 17, 19]. To investigate the putative therapeutic use of CIGB-325 in SARS-CoV-2 infected patients, a Phase I/II clinical trial was conducted [19]. Consistent with the instrumental role of CK2 in SARS-CoV-2 infection, CIGB-325 treatment showed clinical benefit as evidenced by a significant reduction of the pulmonary lesions and lesion’s extent at day 7, suggesting a potential antiviral activity in the infected lung epithelium [19]. However, exploring antiviral activity on *in vitro* coronavirus infection models needs to be accomplished.

Here, we explored the putative antiviral effect of CIGB-325 using a bovine coronavirus infection model and further interrogated those molecular events that might support such antiviral effect. First of all, CIGB-325 exhibited a dose-dependent antiviral activity against BCoV-Mebus according to two different experimental readouts, inhibition of the cytopathic effect and reduction of the viral titer assessed by plaque assay in MDBK cells. Similar to the inhibition of SARS-Cov2 infection by CX-4945 previously reported on Vero-E6 cells [12], the treatment with this specific CK2 inhibitor also elicited anti-BCoV-Mebus activity on MDBK cells in this experimental setting. Thus, the anti-CoV activity of two different CK2 inhibitors supports an instrumental role of the CK2-mediated phosphorylation during infection by this type of *Coronaviridae*. In line with the antiviral activity of CIGB-325 in BCoV-Mebus-infected MDBK cells, the levels of BCoV N mRNA were down-regulated by CIGB-325 treatment as determined by qRT-PCR. However, whether this down-regulation is caused by direct inhibition of the viral RNA transcription derived from CK2 blocking or it is rather a consequence of the antiviral activity by CIGB-325, remains to be elucidated. Accordingly, we also observed a reduction in BCoV M and N protein levels by western blot and *in situ* immunocytochemistry, suggesting an impairment of the viral protein synthesis machinery by CIGB-325.

As CIGB-325 impairs the CK2-mediated phosphorylation through binding to acidic phosphoacceptor domain at the substrates [13], and considering that the coronavirus N protein is a predicted phosphoprotein with multiple putative CK2 phosphorylation sites [33], we also investigated whether CIGB-325 targets the BCoV-N protein. The *in vitro* and *in vivo* pull-down followed by western blot experiments confirmed the typical physic interaction between CIGB-325 and the N protein from BCoV-Mebus. However, additional experiments are required to decipher if CIGB-325 binds directly to BCoV-Mebus N protein or indirectly through unknown cellular protein complexes. Prompted by recent evidences that have shown physical interaction and co-localization between the SARS-CoV-2 N protein and protein kinase CK2 [12, 34], we looked for such interaction in our model. Data from immunoprecipitation experiments clearly corroborated the interaction between CK2 and viral N protein in the context of BCoV-Mebus infection, thus pointing out this protein as a putative CK2 substrate.

Beyond the physical interaction of CIGB-325 with the viral N protein, we wanted to explore whether other viral and host proteins could be targeted by CIGB-325, thus being relevant for its antiviral activity on BCoV-Mebus infection. Interestingly, the CIGB-325 interactomic profile on infected MDBK cells confirmed the interaction between viral N protein with CIGB-325 and also the interaction with NSp2a. Regarding host identified proteins, those representing functional clusters participating in protein folding, cytoskeleton organization and cell cycle were the most representative from the constructed protein-protein interaction network as well as functional enrichment analysis. Interestingly, these three cellular processes are all relevant for the coronavirus life cycle [12, 40, 41].

Furthermore, we interrogated the CIGB-325 interactomic profile for the presence of CK2 substrates and found that 30% (15 proteins) corresponded to previously validated CK2 substrates [31, 32]. A priori, these data reinforce the notion that inhibition of the CK2-mediated phosphorylation could play a central role mediating the CIGB-325 antiviral activity during BCoV-Mebus infection. However, three different CK2 substrates targeted by CIGB-325 might be of particular relevance concerning the antiviral activity on BCoV-Mebus infection due to its already known proviral function. That is the case of B23/NPM protein, a major target for CIGB-325 in solid tumor cells [42], that plays an important role as a proviral chaperone in animal and human virus infections [43–45]. Moreover, the myosin heavy chain 9 (MYL9) is another bonafide CK2 substrate whose phosphorylation at the S1943 is upregulated during infection of Vero-E6 cells by SARS-CoV-2 [12]. Likewise, the heat shock protein 90 alpha family class B member 1 (HSP90AB1) has been identified as a viral RNA interacting protein significantly up-regulated during the SARS-CoV-2 replicative cycle in the RNA-bound proteome on infected lung cells. Consistently, compounds targeting this protein showed a strong inhibition of SARS-CoV-2 protein production [46]. Therefore, B23/NPM, MYH9 and HSP90AB1 represent cellular CK2 substrates whose inhibition by CIGB-325 along with that on viral N protein, may impact the BCoV-Mebus viral machinery eliciting antiviral activity. However, our data do not preclude that other CK2 substrates unveiled among CIGB-325 interacting proteins might be also involved in the antiviral activity.

The CIGB-325 interactomic data originated from different modalities of pull-down experiments here is rather linked to the ability to target the phosphoaceptor domain at the CK2 substrates initially described for this peptide inhibitor [13] However an alternative mechanism of inhibiting the CK2-mediated phosphorylation by direct targeting the CK2α catalytic subunit of the enzyme was recently described for CIGB-325 [15]. Importantly, our *in vivo* pull-down experiments revealed that CIGB-325 can also interact with CK2 catalytic subunit in MDBK cells as it has been previously evidenced in lung cancer and leukemia cells [14, 15]. In line with the abovementioned, the treatment with CIGB-325 showed an impairment of the PI3K/AKT pathway, a CK2 mediated signaling pathway, evaluated by the phosphorylation of downstream signaling protein RPS6. Noteworthy, whether this dual anti-CK2 inhibitory mechanism described for CIGB-325 in other cellular contexts could also be running in parallel, remains to be elucidated.

Further than describing the CIGB-325 antiviral activity on BCoV-Mebus infection and provide clues on the molecular basis of such effect, our data could also suggest an immediate therapeutic tool to treat bovine coronavirus infection. In fact, BCoV is an important livestock pathogen with a high prevalence worldwide causing respiratory disease and diarrhea in calves and winter dysentery in adult cattle with an important loss-inflicting factor in the cattle industry [47].

## 5. Conclusions

In conclusion, we have described for the first time the *in vitro* antiviral activity of CIGB-325 during the BCoV-Mebus infection which corroborates the instrumental role of the CK2-mediated phosphorylation for life cycle of coronavirus. Accordingly, CIGB-325 targeted viral proteins, an array of host CK2 substrates and the CK2 enzyme itself which could all represent crucial molecular clues supporting the antiviral activity. Our data unveil the feasibility of inhibiting CK2 as a promising pharmacological strategy to treat Coronavirus infection and point out CIGB-325 as a putative antiviral drug to treat BCoV and other human Betacoronavirus.

## Author Contributions

Conceptualization, S.E.P., Y.P.; methodology, A.C.R., G.V.P., E.C., D.A., Y.R., A.R.-U., D.V. and V.F.; formal analysis, A.C.R., M.R., D.V. and Y.R.; investigation, A.C.R., G.V.P., M.R., D.V., E.C. and M.P.R.; writing—original draft preparation, A.C.R.; writing—review and editing, S.E.P., M.R., Y.P., D.V.-B; supervision, S.E.P. Y.P. and Y.K.; project administration, S.E.P.

All authors have read and agreed to the published version of the manuscript.

---

**Figure S1.**
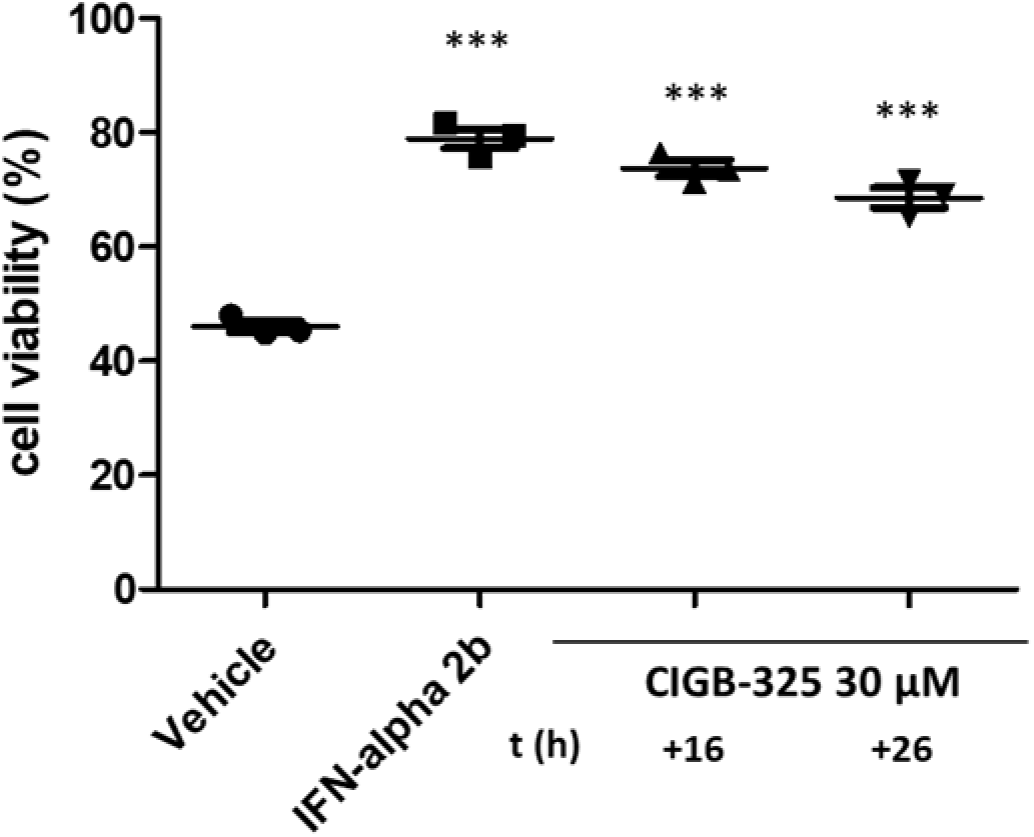
Antiviral activity of CIGB-325 in late stages of BCoV-Mebus infection. MDBK cells were infected with (14 000 TCID50/well), 16 h and 26 h post-infection CIGB-325 (30 μM) was added. Afterward, the cells were incubated for 3 days and the antiviral effect was determined by crystal violet stain. IFN-alpha 2b represents the positive control and was added 1h pre-infection as earlier described. Data is shown as mean ± SD, n = 3. Statistically significant differences between vehicle and drug treatment are represented as \*\**p* < 0.01 and \*\*\**p* < 0.001 determined using one-way ANOVA followed by Dunnett post-test.

**Figure S2.**
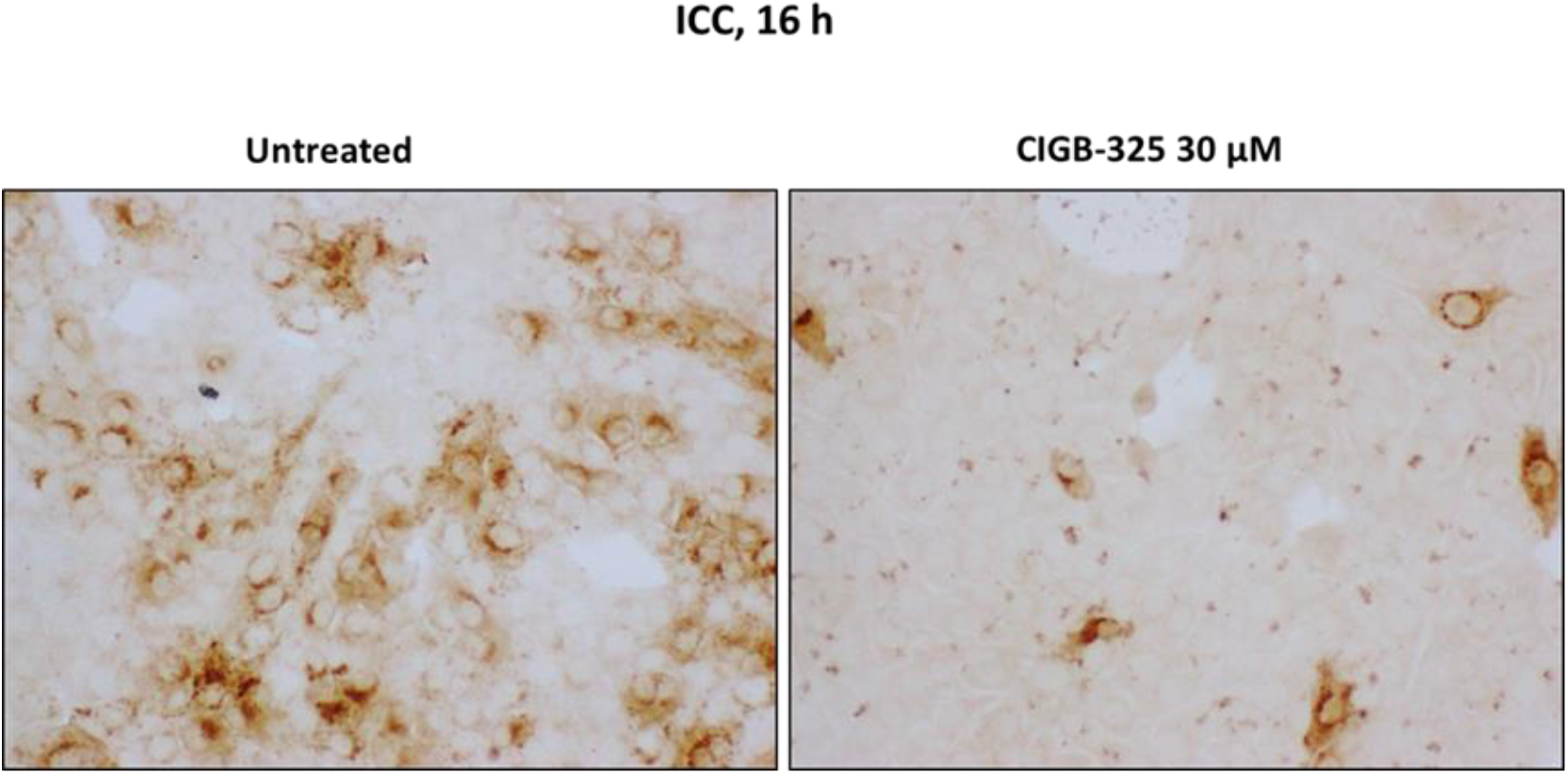
Impact of CIGB-325 over the accumulation of N and M protein by Immunocytochemistry. MDBK cells were treated with the drugs 1 h before the viral challenge. Afterward, cells were infected with BCoV-Mebus (70 000 TCID50/well) and the appropriate drug’s concentration was maintained for 16 h. After the incubation time, cells were fixed and immunostaining using a human anti-SARS-CoV-2 polyclonal antibody from one Covid-19 convalescent patient, followed by peroxidase-conjugated secondary antibody and addition of substrate.

**Figure S3.**
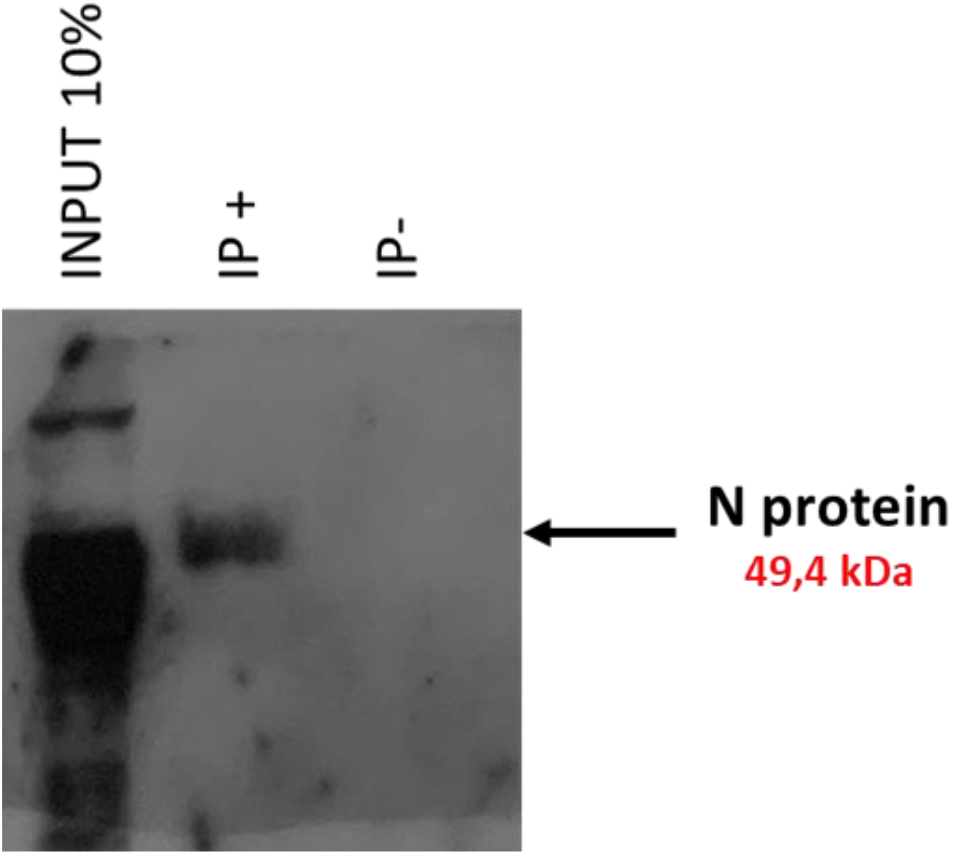
Co-immunoprecipitation of BCoV Mebus N protein and CK2. Cells MDBK were infected 70 000 TCID50. After 48 h post-infection, cell extracts were prepared and immunoprecipitation was performed with anti-CK2α antibody. Cell lysates (lanes marked input) and immunoprecipitate (lanes marked IP) was analyzed on immunoblot with anti-SARS-CoV-2 N protein polyclonal antibody. The position of the N protein is indicated by an arrow. Nonspecific band with anti-N protein antibody is shown.

**Figure S4.**
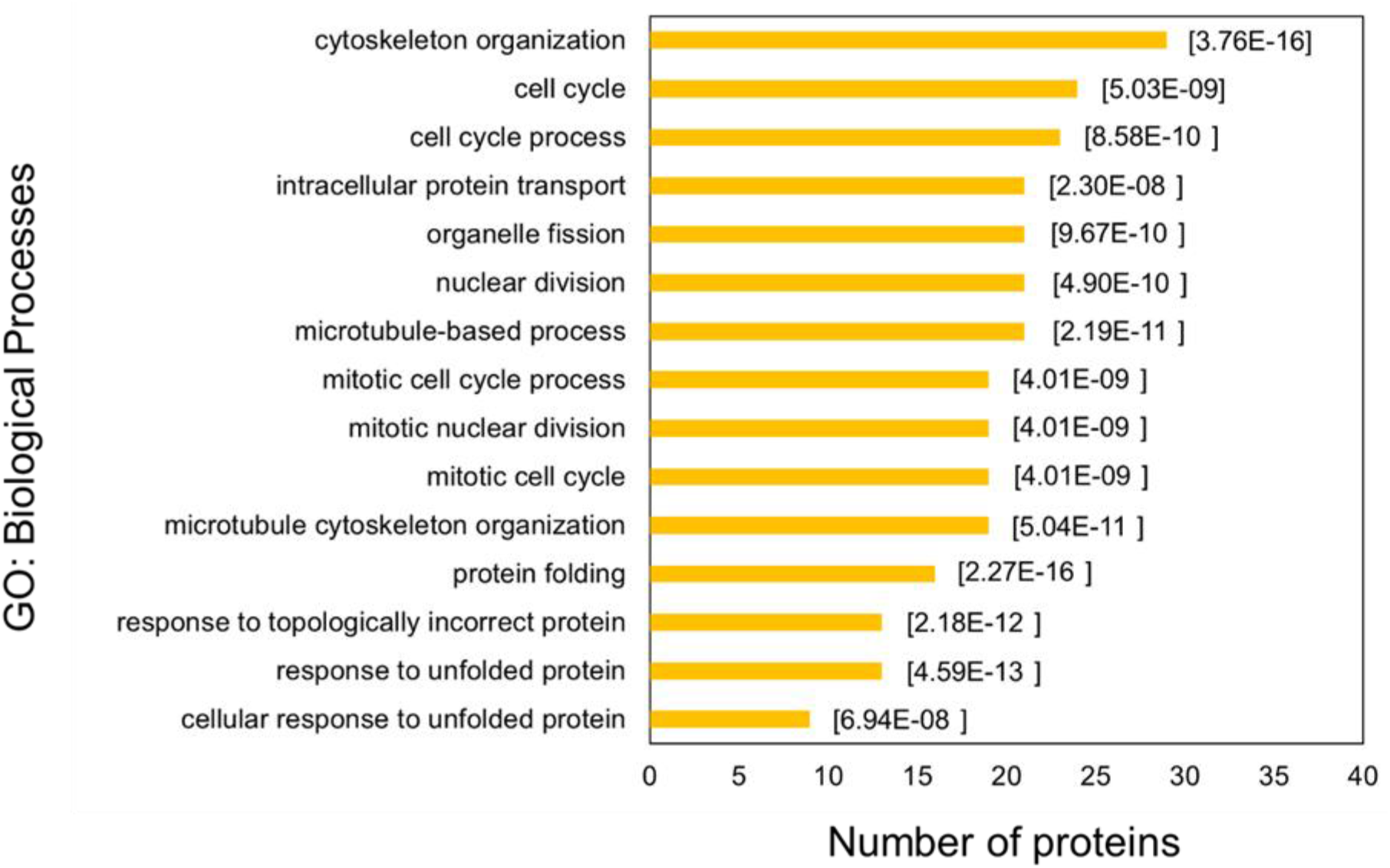
Functional enrichment analysis for CIGB-325 interacting proteins in MDBK infected cells. Biological processes significantly represented in CIGB-325 interactome were identified using annotations from Gene Ontology database. Analysis was performed with ToppFun web-based tool (www.toppgene.cchmc.org/enrichment.jsp/). The *p*-value of each annotation is placed in square brackets.

**Table S1.**
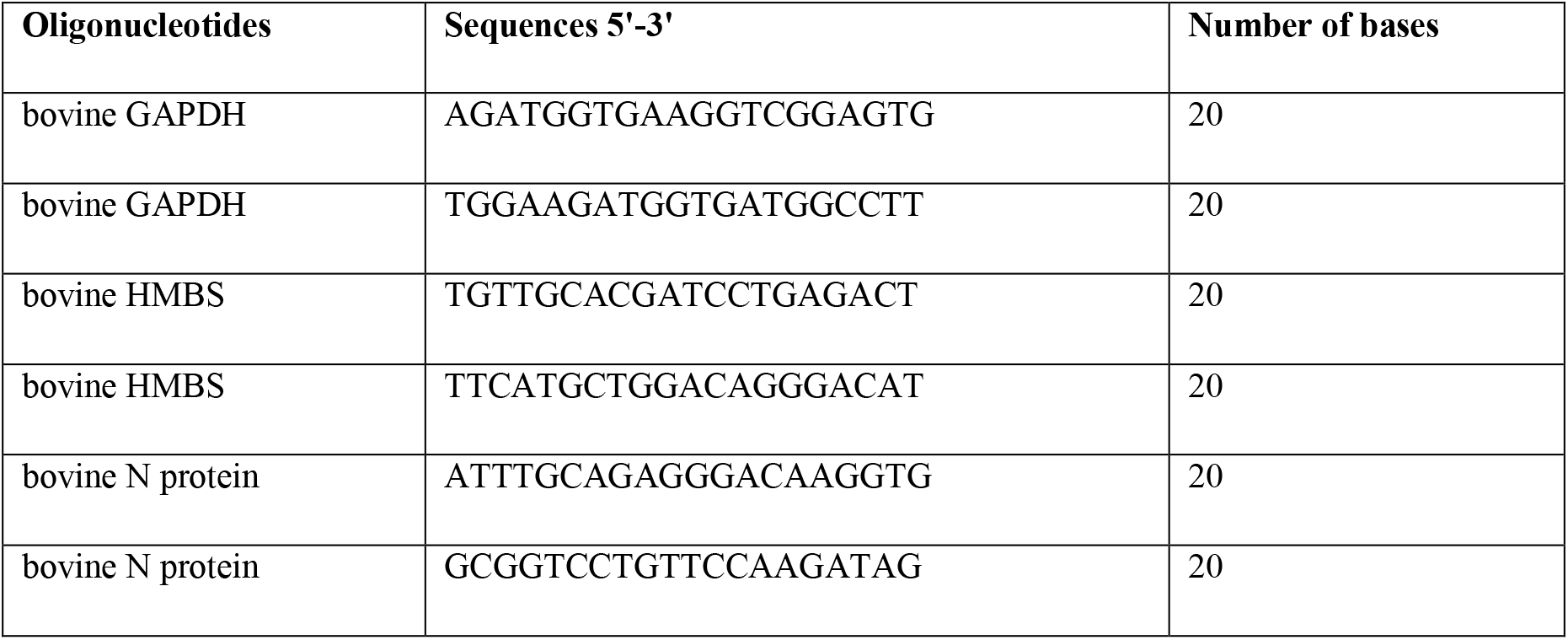
Oligonucleotides sequences.

**Table S2. CIGB-325 interacting proteins**

